# Metaplasticity of cortical glutamatergic LTP by diurnal intracellular chloride dynamics

**DOI:** 10.1101/2023.07.18.549465

**Authors:** Hannah Alfonsa, Vladyslav V. Vyazovskiy, Colin J. Akerman

## Abstract

Neural plasticity varies depending on the time of day and preceding sleep-wake history. It is unclear however, how diurnal changes in cellular physiology modulate a neuron’s propensity to exhibit synaptic plasticity. Recently it has been shown that cortical pyramidal neurons exhibit diurnal changes in their transmembrane chloride gradients, which shift the equilibrium potential for GABAA receptors (EGABAA). Here we demonstrate that diurnal EGABAA affects membrane potential dynamics and glutamatergic long-term potentiation (LTP) elicited by high-frequency spiking activity in pyramidal neurons of mouse cortex. More depolarized EGABAA values associated with the active period facilitate LTP induction by promoting residual depolarization during synaptically-evoked spiking. Diurnal differences in LTP can be reversed by switching the EGABAA-dependent effects on membrane potential dynamics, either by direct current injection or pharmacologically altering EGABAA. These findings identify EGABAA as a metaplastic regulator of glutamatergic synaptic potentiation, which has implications for understanding synaptic plasticity during waking and sleep.

## Introduction

The time of day and preceding sleep-wake history affect the brain’s capacity to process information and are thought to involve modifications in synaptic efficacy ^1,2^. In agreement with this, there has been growing evidence to support sleep-wake regulation of plasticity processes in the brain. For example, studies have shown sleep-wake dependent changes in structural, molecular, and electrophysiological markers of synaptic plasticity ^3–8^. When investigated directly in cortex or hippocampus, the ability to produce long-term potentiation (LTP) at excitatory glutamatergic synapses can vary across the 24h cycle, as a function of sleep-wake history, and depending upon the sleep-wake state ^4,6,8–12^. Other forms of plasticity, such as synaptic scaling, have also been shown to vary with the sleep-wake cycle under normal conditions ^13^, or when there is a homeostatic challenge such as sensory deprivation ^14,15^. Indeed, the regulation of synaptic plasticity is thought to be a major function of sleep ^2,16,17^.

Whilst there is strong evidence for altered plasticity, it is not known how sleep-wake changes in neuronal physiology affect the mechanisms that are responsible for the induction and expression of synaptic plasticity. Recent work has revealed sleep-wake related changes in transmembrane chloride gradients and the equilibrium potential for GABA_A_ receptors (E_GABAA_) in cortex, which are mediated by chloride co-transporter activity ^18^. This work established that recent sleep is associated with more negative E_GABAA_ values in cortical pyramidal neurons, which result in stronger fast synaptic inhibition by hyperpolarizing the postsynaptic membrane. Meanwhile, recent waking is associated with more positive E_GABAA_ values, which favour weaker fast synaptic inhibition and can result in membrane depolarization via the GABA_A_ receptor ^18–21^.

As activity propagates through cortex, feedforward and feedback inhibitory circuits ensure that the glutamatergic synaptic inputs received by a pyramidal neuron, are quickly followed by GABAergic synaptic inputs on a timescale of milliseconds ^22–24^. Changes in E_GABAA_ are therefore predicted to shape membrane potential dynamics during periods of cortical activity ^18,22,24–27^. Here we investigate how sleep-wake history dependent changes in E_GABAA_ affect the postsynaptic membrane potential during synaptically-evoked high frequency spiking activity, which can induce long-term potentiation (LTP) at glutamatergic synapses onto layer 5 (L5) pyramidal neurons ^28^. In the primary somatosensory cortex (S1) of the mouse, we find that depolarizing E_GABAA_ values associated with recent waking promote residual membrane depolarization during synaptically-evoked spike trains. Depolarization between action potentials has been shown to modulate LTP induction via spike-timing-dependent plasticity (STDP) mechanisms ^28–32^ and we reveal that depolarizing E_GABAA_ values facilitate the induction of glutamatergic LTP. In contrast, hyperpolarizing E_GABAA_ values associated with recent sleep reduce membrane depolarization during spike trains and attenuate LTP. Consistent with this being an effect upon LTP induction, varying levels of synaptic potentiation at different times of day can be reversed by counteracting the effects of E_GABAA_ on the membrane potential or by pharmacologically shifting E_GABAA_. These data reveal that E_GABAA_ dynamics are a key regulator of glutamatergic synaptic potentiation across the 24h cycle.

## Results

### E_GABAA_ exhibits diurnal variation in L5 pyramidal neurons

Mice tend to spend the majority of time asleep during the light period and the majority of time awake during the dark period ^33^. In line with recent work ^18^, we assessed E_GABAA_ at two time points within the 24 h cycle - Zeitgeber time 3 (ZT3) when the animals have been recently asleep, and ZT15 when the animals have been recently awake (**Fig. 1a**). Acute slices of primary somatosensory cortex (S1) were prepared at these two time points. Layer 5 (L5) pyramidal neurons were targeted for gramicidin perforated patch clamp recordings, which preserve transmembrane chloride gradients and therefore allow us to study intact GABA_A_ receptor (GABA_A_R) signaling (**Fig. 1a**;^34^). Consistent with previous evidence from multiple areas of cortex ^18^, current clamp recordings in which GABA_A_Rs were activated via a brief, focal somatic puff of GABA, typically elicited hyperpolarizing responses at ZT3, but depolarizing responses at ZT15 (**Fig. 1b**). Ramp protocols conducted in voltage clamp mode confirmed that this was associated with a shift in the equilibrium potential for the GABA_A_R (E_GABAA_), with E_GABAA_ being more depolarizing at ZT15 than at ZT3 (**Fig. 1c**). No difference was observed in the resting membrane potential and, as a result, the GABA_A_R driving force was more depolarizing at ZT15 compared to ZT3 (**Fig. 1d**).

**Fig 1.**
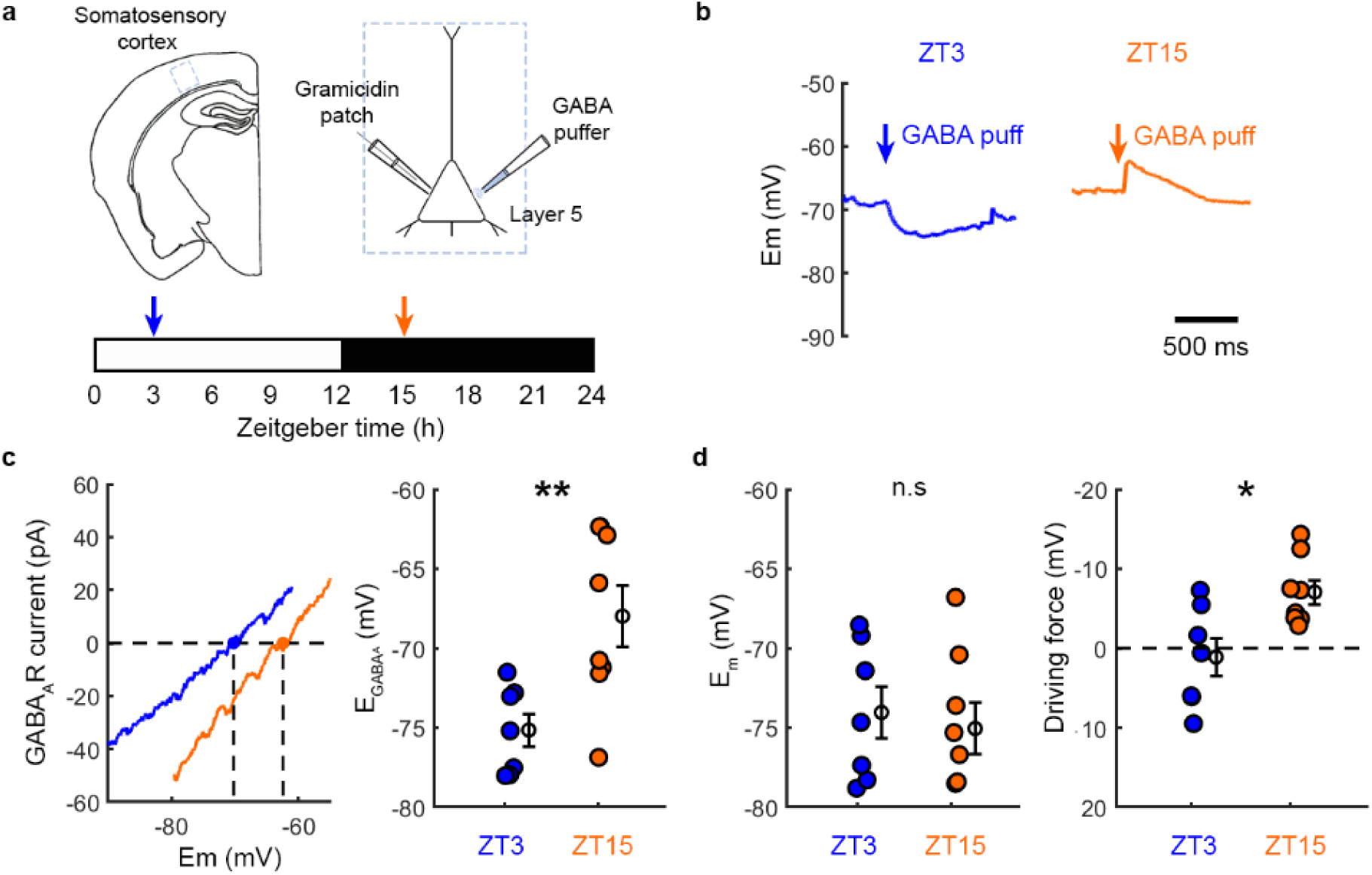
E_GABAA_ exhibits diurnal variation in L5 pyramidal neurons. **a.** Acute brain slices were prepared from primary somatosensory cortex (S1) either at Zeitgeber time 3 (ZT3) during the light period when mice tend to sleep, or at ZT15 during the dark period when mice tend to be awake (indicated by the blue and orange arrows, respectively). Gramicidin perforated patch clamp recordings compared GABA_A_R signalling in layer 5 (L5) pyramidal neurons and GABA_A_Rs were activated by focal delivery of GABA to the neuron’s soma. **b.** Example current clamp (I=0 mode) recordings show the effect of GABA_A_R activation in a slice prepared at ZT3 or ZT15. **c.** GABA_A_R IV curves measured from ramp protocols conducted in voltage clamp mode (left). E_GABAA_ is the membrane potential at which the GABA_A_R current equals zero and was more depolarized at ZT15 compared to ZT3 (right, blue: 7 neurons, 7 slices, 3 animals; orange: 8 neurons, 8 slices, 5 animals; **p=0.0075, unpaired t-test; t=3.16; df=13; d=1.64). **d.** No difference was observed in the resting membrane potential (E_m_; left, blue: 7 neurons, 7 slices, 3 animals; orange: 8 neurons, 8 slices, 5 animals; p=0.67, unpaired t-test; t=0.44; df=13; d=0.23). The GABA_A_R driving force, calculated as E_GABAA_ minus the neuron’s resting membrane potential, was more depolarizing at ZT15 than at ZT3 (right, blue: 7 neurons, 7 slices, 3 animals; orange: 8 neurons, 8 slices, 5 animals; *p=0.011, unpaired t-test; t=2,96; df=13; d=1.53).

### Diurnal variation in E_GABAA_ predicts the degree of membrane depolarization during synaptically-evoked spiking

To investigate how diurnal changes in E_GABAA_ affect membrane potential dynamics and synaptic plasticity, we examined L5 pyramidal neuronal responses to a canonical cortical circuit motif comprising monosynaptic excitation and disynaptic inhibition ^22–24^. To establish the conditions for activating this circuit, whole-cell voltage clamp recordings were initially performed whilst the excitatory-inhibitory input sequence was elicited via a stimulating electrode placed in lower L2/3 (**Fig. 2a**). In slices prepared at both ZT3 and ZT15, single stimuli elicited fast, monosynaptic excitatory postsynaptic currents (EPSCs) when the L5 pyramidal neuron was held at E_GABAA_ (-80 mV under whole cell recording conditions). Whereas the same stimulus elicited inhibitory postsynaptic currents (IPSCs) when the pyramidal neuron was held at the equilibrium potential for glutamate receptors (E_Glu_; 0 mV), and the timing of these IPSCs was consistent with a disynaptic delay (∼ 2 ms; **Fig. 2a**; ^24^).

**Fig 2.**
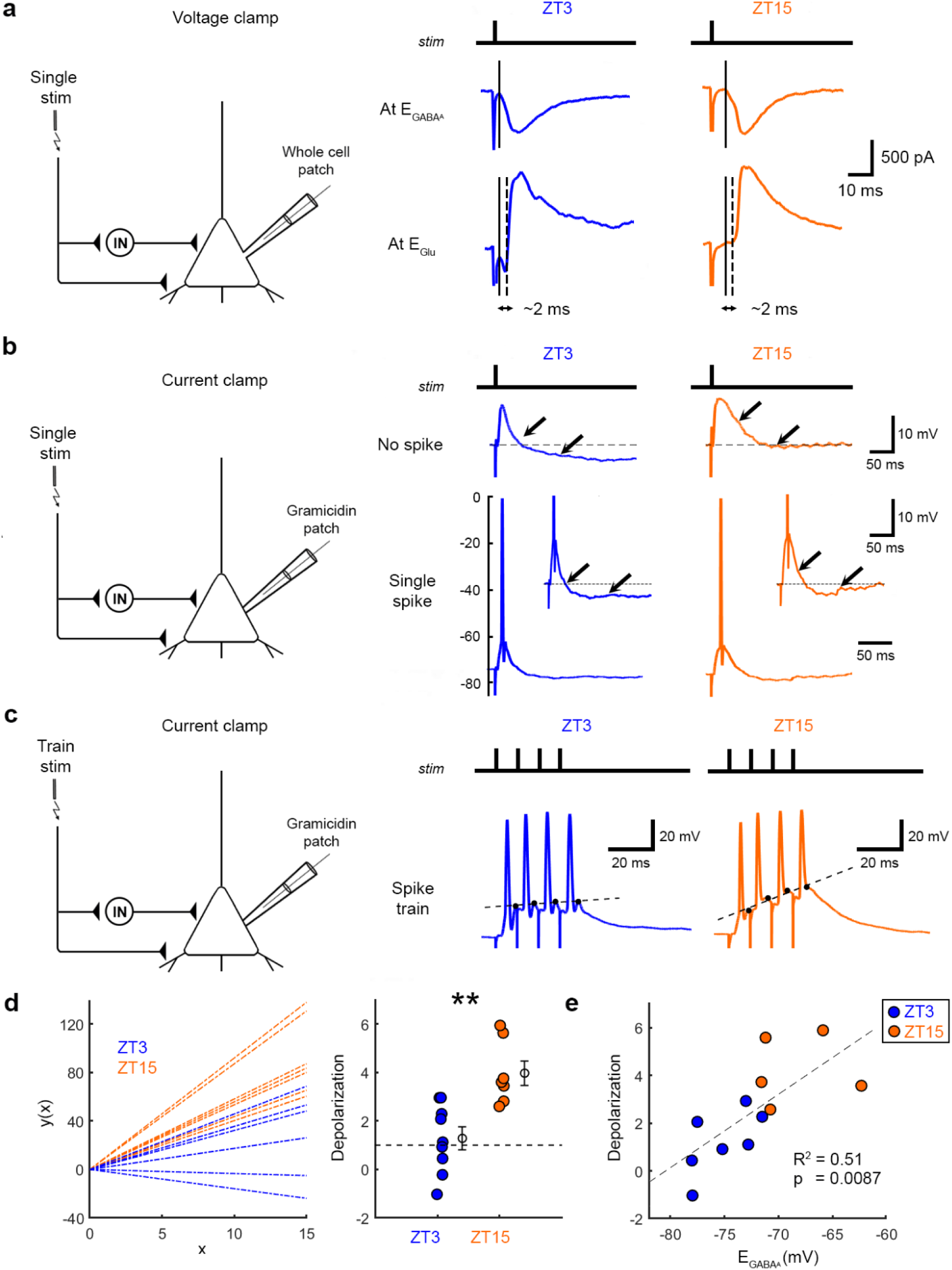
Diurnal variation in E_GABAA_ predicts the degree of membrane depolarization during synaptically-evoked spiking. **a.** Whole-cell patch clamp recordings in voltage clamp mode were performed from L5 pyramidal neurons in S1, whilst an excitatory-inhibitory circuit was activated via a stimulating electrode in lower L2/3 (left). Example recordings (right) for a neuron at ZT3 (blue) and ZT15 (orange) show the monosynaptic EPSC recorded at E_GABAA_ and disynaptic IPSC recorded at E_Glu_, following a single stimulus. **b.** Gramicidin perforated patch clamp recordings in current clamp mode (I=0) were used to monitor membrane potential responses to a single stimulus (left). Example recordings (right) for a neuron at ZT3 and ZT15 when the stimulus intensity was set either below the spike threshold (No spike) or above the spike threshold (Single spike). Arrows and insets highlight the effects of the disynaptic IPSP on the subthreshold and after-spike membrane potential. **c.** Same arrangement as in ‘b’, used to monitor responses to a high frequency stimulus train (left; 100 Hz, 4 pulses). Example recordings (right) showing high frequency synaptically-evoked spiking (‘Spike train) elicited in a neuron at ZT3 and at ZT15. The neuron at ZT15 exhibits greater residual depolarization between action potentials of the spike train. This depolarization was defined as the slope of the linear fit to the post-spike depolarization (dashed lines). **d.** Population data showing linear fits to the residual depolarization (left). Depolarization was greater at ZT15 when E_GABAA_ is more depolarized (right, blue: 9 neurons, 9 slices, 7 animals; orange: 7 neurons, 7 slices, 5 animals; **p=0.0016, unpaired t-test; t=3.91; df=14; d=1.97). **e.** E_GABAA_ showed a positive linear correlation with the degree of residual depolarization (12 neurons, 12 slices, 8 animals; r=0.72; p=0.0087).

Subsequent experiments used gramicidin perforated patch clamp recordings to preserve the L5 pyramidal neuron’s native E_GABAA_ (**Fig. 2b**). Under these conditions, current clamp recordings revealed differences between ZT3 and ZT15 in terms of the L5 neuron’s excitatory-inhibitory synaptic responses. Consistent with their more hyperpolarized E_GABAA_ (**Fig. 1**) ^18^, L5 neurons at ZT3 tended to exhibit more hyperpolarized potentials during the disynaptic inhibitory window following stimulation. This was apparent when the stimulus intensity was set such that the L5 pyramidal neuron did not reach action potential threshold (‘no spike’), or when the stimulus intensity was increased to a suprathreshold level that elicited an action potential in the L5 pyramidal neuron (‘single spike’; **Fig. 2b**). In contrast, and consistent with their more depolarized E_GABAA_ (**Fig. 1**), L5 neurons at ZT15 tended to remain more depolarized and showed less hyperpolarization during the disynaptic inhibitory window (**Fig. 2b**).

These differences between ZT3 and ZT15 were also evident when the suprathreshold stimulus was used to evoke trains of postsynaptic spiking activity in the L5 pyramidal neuron (**Fig. 2c**). High frequency trains of suprathreshold stimulation (100 Hz, 4 stimuli, L5 spike with each stimulus) produced synaptically-evoked trains of action potentials in the L5 pyramidal neuron (‘spike train’). Consistent with their more depolarized E_GABAA_ and its effects during the disynaptic inhibitory window, neurons at ZT15 exhibited greater residual depolarization between action potentials, which increased over the spike train (**Fig. 2c**). This depolarization could be quantified as the slope of the linear fit to the post-spike depolarization (**Fig. 2c**), was found to be significantly greater at ZT15 than at ZT3 (**Fig. 2d**), and was positively correlated with the neuron’s E_GABAA_ (**Fig. 2e**). These data reveal that changes in E_GABAA_ shape membrane potential dynamics elicited by excitatory-inhibitory input sequences onto L5 pyramidal neurons, such that more depolarized E_GABAA_ values following recent waking are associated with greater residual depolarization during synaptically-evoked spiking activity.

### L5 pyramidal neurons exhibit diurnal differences in glutamatergic LTP

High frequency presynaptic and postsynaptic spiking activity has been shown to elicit LTP in L5 pyramidal neurons ^28^. To examine whether such forms of LTP vary between ZT3 and ZT15, we prepared acute cortical brain slices at these time points, and performed gramicidin perforated patch clamp recordings from L5 pyramidal neurons (**Fig. 3a**). Monosynaptic EPSP peak amplitude was monitored in response to electrical stimulation of presynaptic input from L2/3 and compared before and after an LTP induction protocol, which involved eliciting trains of high frequency synaptically-evoked postsynaptic spikes (four spikes at 100 Hz, as in **Fig. 2c**), repeated 100 times at an interval of 10 s (**Fig. 3b**). This protocol resulted in greater LTP at ZT15 than at ZT3 (**Fig. 3c-d**). These differences in LTP levels could not be accounted for by the postsynaptic spiking activity during the induction protocol (**Fig. 3e**), but did exhibit a positive correlation with the degree of residual depolarization that the neuron exhibited during the induction protocol (**Fig. 3f**). In addition, the L5 neuron’s E_GABAA_ was positively correlated with the level of LTP that the neuron exhibited (**Fig. 3g**). Combined, these data suggest a link between diurnal variation in E_GABAA_, membrane depolarization, and the level of LTP induced by high frequency presynaptic and postsynaptic activity.

**Fig 3.**
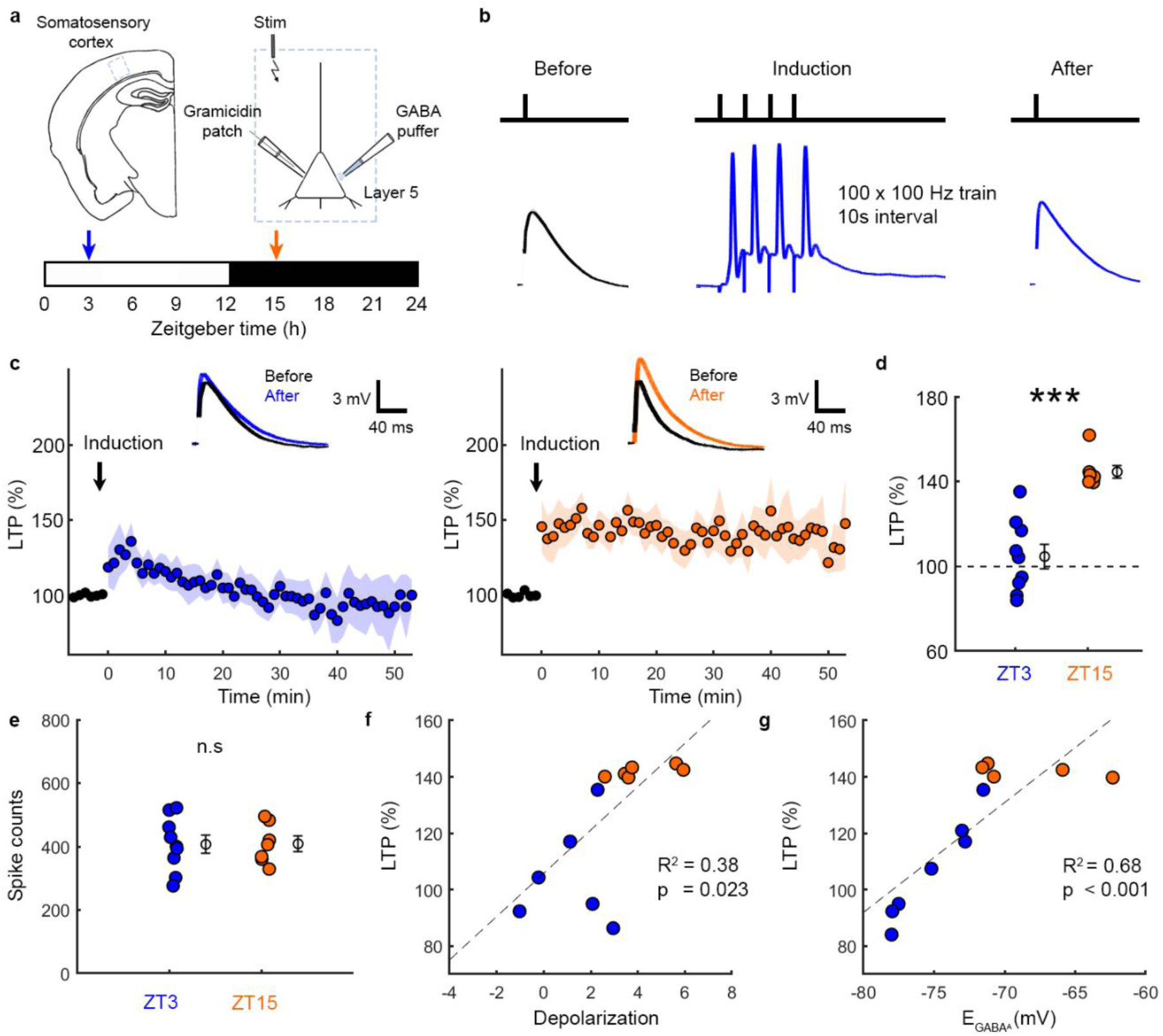
L5 pyramidal neurons exhibit diurnal differences in glutamatergic LTP. **a.** Acute brain slices were prepared at ZT3 or ZT15. An input pathway was stimulated whilst a gramicidin perforated patch clamp recording was performed from a L5 pyramidal neuron in S1. **b.** EPSP amplitude was monitored before and after the LTP induction protocol. LTP induction involved eliciting trains of high frequency synaptically-evoked postsynaptic spikes (four spikes at 100 Hz), repeated 100 times at an interval of 10 s. **c.** Population data showing the percentage LTP (EPSP peak amplitude, normalized to the mean baseline) before and after LTP induction (arrow) for the ZT3 (left) and ZT15 (right) condition. Inset shows example mean EPSPs before and after LTP induction. **d.** LTP was larger at ZT15 compared to ZT3 (blue: 9 neurons, 9 slices, 7 animals; orange: 7 neurons, 7 slices, 5 animals; ***p=0.0002, Mann-Whitney test; d=2.86). **e.** No difference was observed in the spike counts during LTP induction (blue: 9 neurons, 9 slices, 7 animals; orange: 7 neurons, 7 slices, 5 animals; p=0.965, unpaired t-test; t=0.044; df=14; d=0.02). **f.** The residual depolarization between postsynaptic spikes showed a positive linear correlation with the amount of LTP (13 neurons, 13 slices, 11 animals; r=0.62; p=0.023). **g.** A neuron’s E_GABAA_ showed a positive linear correlation with the amount of LTP the neuron exhibited (12 neurons, 12 slices, 8 animals; r=0.82; p<0.001). Data represent mean ± sem.

### Postsynaptic membrane potential determines diurnal differences in LTP induction

Previous work has identified postsynaptic membrane potential as a critical determinant of LTP induction at L5 glutamatergic synapses, revealing that LTP can be gated according to the degree of membrane depolarization between postsynaptic spikes ^28,29,31,32^. We hypothesized that the diurnal differences observed in LTP at L5 glutamatergic synapses are influenced by the degree of postsynaptic depolarization, which is regulated through the effects of E_GABAA_ upon disynaptic inhibition. To test the first element of this hypothesis, we investigated whether the amount of LTP was influenced by the degree of residual postsynaptic depolarization during synaptically-evoked spike trains. We examined whether a lack of membrane depolarization limits LTP induction at ZT3, by injecting brief depolarizing current pulses (4 ms duration) between the synaptically-evoked postsynaptic spikes (**Fig. 4a**), which successfully increased the L5 pyramidal neuron’s depolarization during the LTP induction protocol (**Fig. 4b-c**). Meanwhile, to address whether membrane depolarization is responsible for the enhanced LTP induction at ZT15, we injected brief hyperpolarizing current pulses (4 ms duration) between the synaptically-evoked postsynaptic spikes (**Fig. 4a**), which successfully decreased the residual depolarization during the LTP induction protocol (**Fig. 4b-c**). Under both of these conditions, we found a striking reversal in the amount of LTP that was elicited. Pairing depolarizing current pulses with the synaptically-evoked spiking protocol enhanced the induction of LTP at ZT3 (ZT3+Depol) compared to control recordings (ZT3; **Fig. 4d-e**). In contrast, pairing hyperpolarizing current pulses with the synaptically-evoked spiking protocol reduced the amount of LTP that was elicited at ZT15 (ZT15+Hyperpol) compared to control recordings (ZT15; **Fig. 4d-e**). Therefore, the diurnal differences in the level of LTP are determined by the amount of residual depolarization between postsynaptic action potentials during induction.

**Fig 4.**
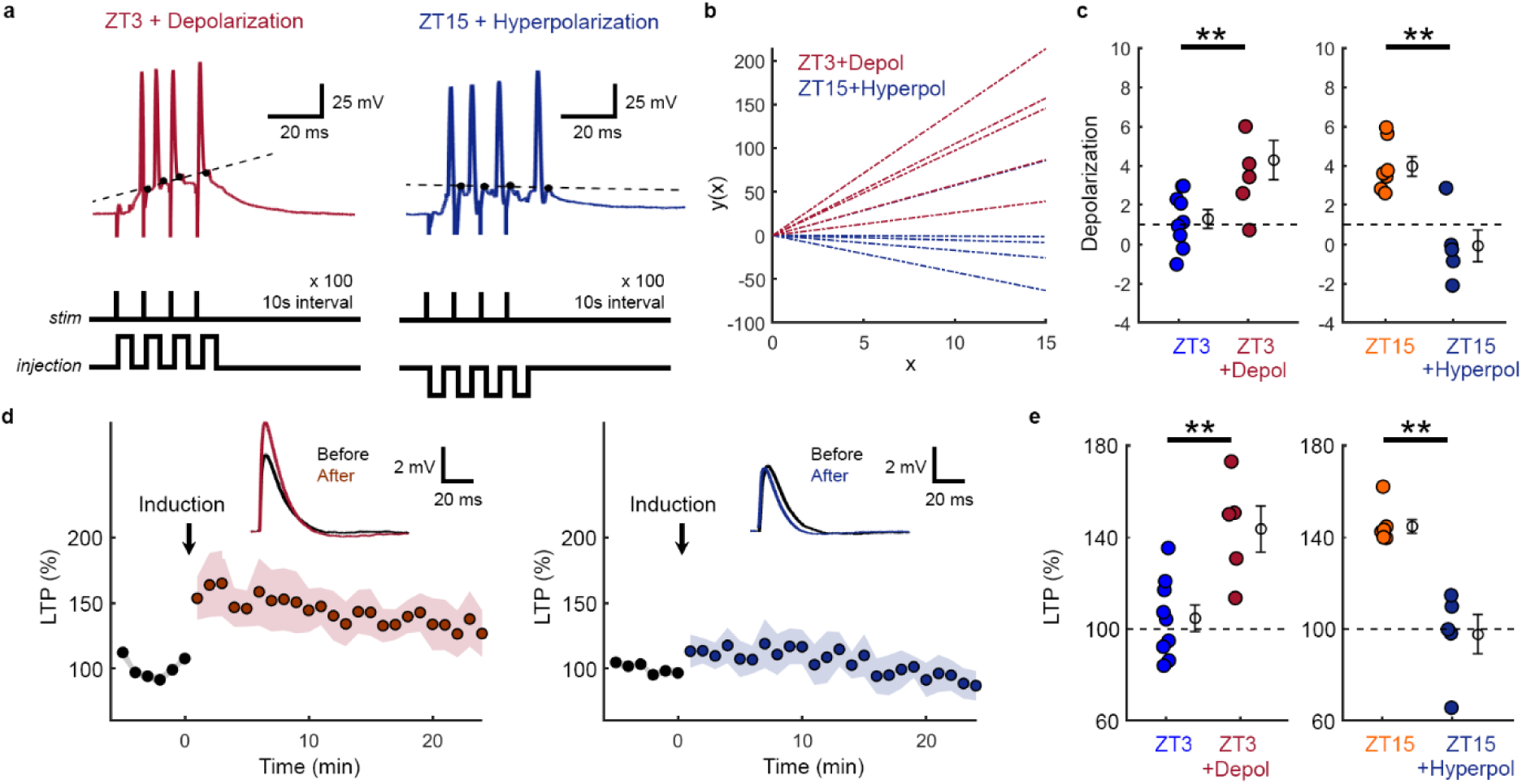
Postsynaptic membrane potential determines diurnal differences in LTP induction. **a.** Gramicidin perforated patch clamp recordings were used to monitor synaptically-evoked spike trains in L5 pyramidal neurons, and brief somatic current pulses (4 ms duration) were delivered to modulate the membrane potential between postsynaptic spikes. Depolarizing current pulses were used to increase depolarization at ZT3 (maroon), whilst hyperpolarizing current pulses were used to decrease depolarization at ZT15 (navy). The resulting residual depolarization was defined as the slope of the linear fit to the post-spike depolarization. **b.** Population data showing linear fits to the residual depolarization. **c.** Depolarization was increased in the ZT3+Depolarization condition compared to control ZT3 (blue: same data as Fig. 2d; maroon: 5 neurons, 5 slices, 5 animals; **p=0.009, unpaired t-test; t=3.109; df=12; d=1.73). Depolarization was reduced in the ZT15+Hyperpolarization condition compared to control ZT15 (orange: same data as Fig. 2d; navy: 5 neurons, 5 slices, 3 animals; **p=0.0011, unpaired t-test; t=4.5; df=10; d=2.64). **d.** Population data showing the percentage LTP (EPSP peak amplitude, normalized to the mean baseline) before and after induction (arrow) with either the ZT3+Depolarization protocol (left) or ZT15+Hyperpolarization protocol (right). The LTP induction protocols involved eliciting trains of high frequency synaptically-evoked postsynaptic spikes (four spikes at 100 Hz), combined with either depolarizing or hyperpolarizing current pulses as shown in ‘a’, and repeated 100 times with 10 s intervals between trains. Inset shows example mean EPSPs before and after the LTP induction. **e.** LTP was enhanced at ZT3 by increasing the post-spike depolarization during induction (blue: same data as Fig. 3d; maroon: 5 neurons, 5 slices, 5 animals; **p=0.0034, unpaired t-test; t=3.64; df=12; d=2.03). Whilst LTP was attenuated at ZT15 by reducing the post-spike depolarization during induction (orange: same data as Fig. 3d; navy: 5 neurons, 5 slices, 3 animals; **p=0.0025, Mann-Whitney test; d=3.48). Data represent mean ± sem.

### Diurnal variation in EGABA_A_ gates LTP induction

The second element of our hypothesis regarding diurnal differences in L5 LTP, was that the effects are mediated via the influence of E_GABAA_ upon the membrane potential. Consistent with this, we had already observed positive correlations between E_GABAA_ and both the degree of depolarization during synaptically-evoked spiking activity (**Fig. 2**) and the amount of LTP at ZT3 and ZT15 (**Fig. 3**). To test this aspect of the hypothesis directly, we used chloride co-transporter blockers to either produce a depolarizing shift in E_GABAA_ at ZT3 with an antagonist of the chloride extruder, KCC2 (VU0463271, 10 μm), or a hyperpolarizing shift in E_GABAA_ at ZT15 with an antagonist of the chloride intruder, NKCC1 (bumetanide, 10 μm) ^18,35–37^. Under these conditions, we then explored the effects upon LTP induction. Blocking KCC2 to cause a depolarizing shift in E_GABAA_ at ZT3 (ZT3+VU) increased the L5 pyramidal neuron’s residual depolarization during the synaptically-evoked spike trains of the LTP induction protocol (**Fig. 5a-c**). Meanwhile blocking NKCC1 to cause a hyperpolarizing shift in E_GABAA_ at ZT15 (ZT15+Bume) resulted in reduced residual depolarization during the LTP induction protocol (**Fig. 5a-c**).

**Fig 5.**
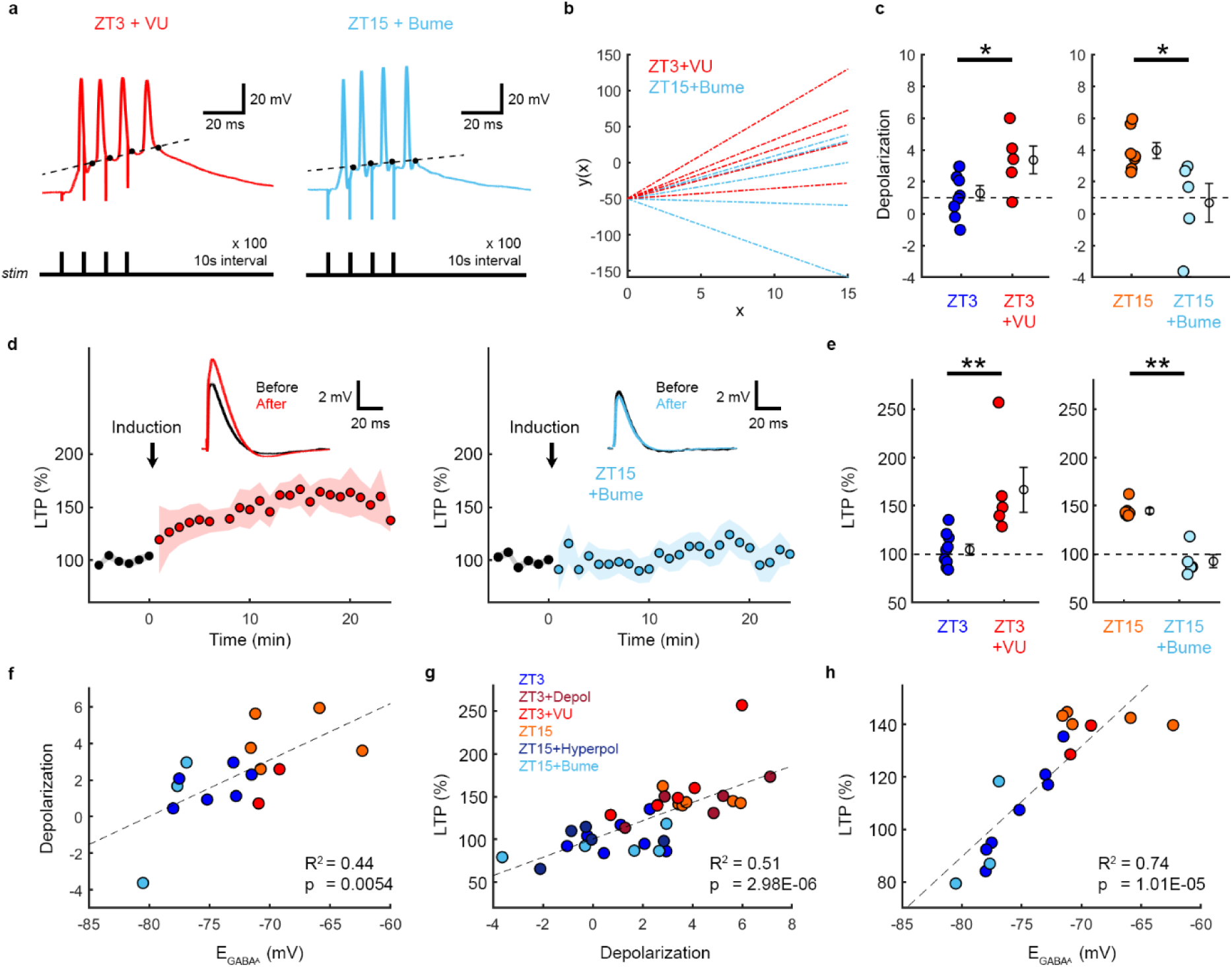
Diurnal variation in EGABA_A_ gates LTP induction. **a.** Gramicidin perforated patch clamp recordings were used to monitor synaptically-evoked spiking activity in L5 pyramidal neurons, in the presence of a KCC2 blocker (VU) at ZT3 (red), or an NKCC1 blocker (Bume) at ZT15 (light blue). **b.** Population data showing linear fits to the residual depolarization. **c.** Depolarization was greater in the ZT3+VU condition compared to control ZT3 (blue: same data as Fig. 2d; red: 5 neurons, 5 slices, 4 animals; *p=0.0166, unpaired t-test; t=2.782; df=12; d=1.55). Depolarization was lower in ZT15+Bume condition compared to control ZT15 (orange: same data as Fig. 2d; light blue: 5 neurons, 5 slices, 4 animals; *p=0.0183, unpaired t-test; t=2.816; df=10; d=1.65). **d.** Population data showing the percentage LTP (EPSP peak amplitude, normalized to the mean baseline) before and after induction (arrow) in the ZT3+VU condition (left) and ZT15+Bume (right) condition. Inset shows example mean EPSPs before and after the LTP induction protocols. **e.** LTP was enhanced at ZT3 when induced in the presence of VU (blue: same data as Fig. 3d; red: 5 neurons, 5 slices, 4 animals; **p=0.002, Mann-Whitney test; d=1.88). Whilst LTP was attenuated at ZT15 when induced in the presence of bumetanide (orange: same data as Fig. 3d; light blue: 5 neurons, 5 slices, 4 animals; **p=0.0025, Mann-Whitney test; d=4.62). **f.** Across all recording conditions, a neuron’s E_GABAA_ showed a positive linear correlation with the degree of residual depolarization that the neuron exhibited during the synaptically-evoked spike trains (16 neurons, 16 slices, 12 animals; r=0.66; p=0.0054). **g.** A neuron’s residual depolarization during the LTP induction protocol showed a positive linear correlation with the amount of LTP that the neuron exhibited (34 neurons, 34 slices, 27 animals; r=0.71; p<0.001). **h.** A neuron’s E_GABAA_ showed a positive linear correlation with the level of LTP (17 neurons, 17 slices, 13 animals; r=0.86; p<0.001). Data represent mean ± sem.

Both of these manipulations reversed the amount of LTP that was elicited. Blocking KCC2 at ZT3 resulted in increased LTP compared to control recordings at the same time of day (**Fig. 5d-e**). Whilst blocking NKCC1 at ZT15 resulted in reduced LTP compared to control recordings at the same time of day (**Fig. 5d-e**). Finally, to assess the relationship between E_GABAA_, residual depolarization and the amount of LTP, we pooled the data collected across all of the experimental conditions. This revealed strong positive correlations between a L5 pyramidal neuron’s E_GABAA_ and the depolarization that the neuron exhibited during synaptically-evoked spiking activity (**Fig. 5f**), between depolarization and the amount of LTP (**Fig. 5g**), and also between E_GABAA_ and the amount of LTP (**Fig. 5h**). Collectively, these data support the conclusion that diurnal differences in E_GABAA_ modulate the induction of LTP via E_GABAA_-dependent effects upon the postsynaptic neuron’s membrane potential dynamics.

## Discussion

The ability to produce long-term potentiation (LTP) at excitatory glutamatergic synapses can vary across the 24h cycle, as a function of sleep-wake history, and depending upon the sleep-wake state ^3–12^. Yet it is not known how LTP is affected by sleep-wake related changes in cellular physiology. Here we reveal that diurnal changes in glutamatergic LTP relate to underlying changes in the equilibrium potential for the GABA_A_ receptor (E_GABAA_), which determine the nature of postsynaptic inhibition ^20,21^. When a L5 pyramidal neuron’s spiking activity is generated by excitatory and inhibitory circuits, more depolarized E_GABAA_ values at ZT15 can result in greater residual depolarization between postsynaptic action potentials, compared to when E_GABAA_ is more hyperpolarized at ZT3. This E_GABAA_–dependent modulation of the membrane potential gates glutamatergic synaptic potentiation. First, the degree of E_GABAA_–dependent residual depolarization correlates with the degree of glutamatergic LTP. Second, directly counteracting the effects of E_GABAA_ upon the membrane potential results in a reversal in the ability to induce glutamatergic LTP at different times of day. And third, shifting E_GABAA_ to have more depolarized values at ZT3, or more hyperpolarized at ZT15, reverses both the level of residual depolarization and LTP at each time point. Thus, we identify diurnal variations in E_GABAA_ as a key factor underlying the induction of glutamatergic LTP in cortex, and a potential target for modulating sleep-wake related changes in the brain’s capacity for plasticity.

These findings extend our previous work ^18^, in which we established that diurnal variations in a cortical pyramidal neuron’s E_GABAA_ reflect the animal’s sleep-wake history. More depolarized E_GABAA_ values are associated with recent wakefulness, altered contributions by chloride cotransporter proteins, and can account for markers of sleep pressure within cortex ^18,38^. The current work establishes for the first time that E_GABAA_ dynamics across the sleep-wake cycle affect the potentiation of glutamatergic cortical synapses. Differences in the amount of glutamatergic LTP can reflect induction mechanisms, expression mechanisms, or a combination of both. As E_GABAA_ alters the pyramidal neuron’s membrane potential dynamics, we interpret our findings as consistent with an effect upon LTP induction, whereby the threshold for inducing LTP is effectively raised or lowered as a function of E_GABAA_. This is supported by the fact that reversing the effects of E_GABAA_ was sufficient to reverse the amount of LTP elicited at both ZT3 and ZT15, in a bidirectional manner. Thus, even though there are sleep-wake related molecular changes at glutamatergic synapses ^7,39^, L5 pyramidal neurons have the potential to express LTP at different points during the sleep-wake cycle. In addition to the effects of E_GABAA_, other diurnal aspects of inhibitory synaptic transmission could contribute to LTP regulation. For instance, the inhibitory synaptic conductances received by pyramidal neuronal populations can increase during the sleep period ^40,41^, which would complement hyperpolarizing shifts in E_GABAA_ to achieve stronger postsynaptic inhibition.

A defining feature of LTP is that its induction exhibits cooperativity ^42^. Traditionally, this has been considered in terms of the excitatory inputs that must combine to satisfy the induction rules in the postsynaptic neuron, such as achieving sufficient depolarization to relieve the voltage-dependent magnesium block of NMDAR receptors (NMDARs) and allow calcium influx ^43,44^. Previous seminal work established a form of cooperativity that gates LTP induction and depends upon the degree of residual membrane depolarization during postsynaptic spiking, when paired with pre-synaptic activity ^28^. It was shown in L5 pyramidal neurons that residual depolarization permits STDP-induced LTP elicited by high frequency spiking trains that satisfy a pre-before-post rule. Considering the NMDAR as a coincidence detector ^44,45^, the residual depolarization is well-placed to reduce the magnesium block and prepare the dendrite for successful backpropagation of a subsequent action potential ^28–32^. Our findings closely align with these observations, but highlight that cooperativity in LTP induction should be considered in terms of both the glutamatergic and GABAergic synaptic inputs received by a neuron ^28,42,44^ and identify E_GABAA_ dynamics as a key parameter determining the cooperation between glutamatergic and GABAergic inputs. The influence of E_GABAA_ upon the neuron’s membrane potential dynamics would be well-placed to regulate the voltage-dependent block of NMDARs. This could be through direct effects of GABA_A_R activation on the membrane potential, as has been seen when E_GABAA_ is depolarizing during development ^25,46^. Alternatively, the influence of E_GABAA_ upon sustained depolarization could affect NMDARs via indirect effects on the degree to which the action potential backpropagates ^47^, given that membrane depolarization has been shown to favour spike backpropagation, most likely through sodium channel activation mechanisms in L5 pyramidal neurons ^28,47^.

It is well established that synaptic plasticity varies with sleep and wakefulness ^3,4,8,48^. Our data show that LTP in L5 pyramidal neurons is greater at times associated with recent wakefulness, when E_GABAA_ is relatively depolarizing. This is consistent with multiple lines of evidence. For example, wakefulness has been associated with an increase in the axon-spine interface ^3^, a structural correlate of synaptic strength. Additionally, wakefulness has been associated with an increase in AMPA receptor levels and their phosphorylation ^4,49^, which are considered molecular correlates of synaptic strength. Electrophysiological measures, such as increased slope and amplitude of evoked cortical responses, and the frequency of miniature excitatory postsynaptic currents, have also been observed with wakefulness ^4,13,40^. Collectively this evidence supports the first aspect of the synaptic homeostasis hypothesis of sleep (SHY) ^16^, which proposes that during wakefulness, the intake and encoding of information leads to synaptic potentiation. Our findings suggest that the wake-associated depolarizing shift in E_GABAA_ forms part of a mechanism that facilitates the induction of synaptic potentiation with wakefulness. The SHY also proposes that synaptic potentiation during wakefulness is linked to the homeostatic regulation of slow-wave activity (SWA) during non-rapid eye movement (NREM) sleep ^16^. In our previous study, we demonstrated that wake-related depolarizing shifts in E_GABAA_ occur locally within cortex, are activity-dependent, and regulate the level of local SWA ^18^. E_GABAA_ regulation could therefore serve as a cell-specific mechanism that links wakefulness, synaptic potentiation and homeostatic regulation of SWA. Given that active cortical regions experience the largest depolarizing shifts in E_GABAA_ ^18^, one prediction is that these should show the greatest synaptic potentiation, as has been shown for SWA levels.

While there is evidence that links synaptic potentiation and wakefulness, previous work has also shown that LTP occurs during sleep and that this plays a key role in consolidating learning and memory processes ^2,8,9^. It is therefore interesting to ask whether E_GABAA_ dynamics could contribute to the consolidation of synaptic changes. Consistent with this possibility, the effects of depolarizing E_GABAA_ upon cortical activity can be observed following the onset of sleep ^18^, suggesting that E_GABAA_ is relatively slow to recover with a change in vigilance state, and could facilitate consolidation processes by lowering the threshold for LTP during early sleep. In this sense, E_GABAA_ could be considered part of a potential ‘priming’ mechanism, as has been proposed ^17^, with the activity-dependent depolarizing E_GABAA_ shifts during waking serving to promote early LTP on a timescale of hours, whilst also priming a neuron so that its synaptic connections might be consolidated during subsequent sleep.

In summary, sleep-wake related changes in a cortical pyramidal neuron’s E_GABAA_ gates the induction of glutamaterigic LTP. These effects represent a form of excitatory-inhibitory cooperativity, whereby depolarizing E_GABAA_ promotes residual depolarization during synaptically-evoked spiking, which facilitates LTP induction. Counteracting these E_GABAA_–dependent membrane potential dynamics, either directly or by pharmacologically targeting chloride cotransporter proteins, can invert the diurnal differences in synaptic plasticity, such that LTP becomes either easier or harder to induce at different points in the sleep-wake cycle. Thus, E_GABAA_ dynamics represent a potential target for modulating diurnal plasticity, and provide a functional link between sleep-wake related changes in a neuron’s physiology, and the mechanisms that are responsible for the induction of synaptic plasticity.

## Acknowledgments

We thank the Akerman lab for advice and comments.

## Author contributions

HA and CJA conceptualized the project and designed the experiments. VV assisted with the interpretation of the data. HA performed and analysed the experiments. HA and CJA wrote the manuscript with advice and input from VV.

## Data availability

Source data are provided with this manuscript.

## Code availability

Custom-made MATLAB code used for analysis is available from the corresponding authors upon reasonable request.

## Methods

### Ethical approval

All experiments were performed in accordance with the United Kingdom Animal Scientific Procedures Act 1986 under personal and project licences granted by the United Kingdom Home Office. Approval was also provided by an Ethical Review Panel at the University of Oxford.

### Animal husbandry

Experiments were performed on male C57BL/6 wild-type mice purchased from Charles River. Animals were maintained under a 12-h:12-h light-dark (LD) cycle. Ambient room temperature was maintained at 22 ± 2 ^о^C and humidity at 50 ± 20%. Animals were aged 4-8 weeks and animal numbers are provided in the figure legends for each experiment. Animals were maintained under a 12-h:12-h light-dark (LD) cycle.

### Acute brain slices

Acute cortical brain slices were prepared for electrophysiological recordings at ZT3 or ZT15. Animals from the same litter were randomly assigned to the different ZT conditions. Animals were collected, and immediately sacrificed by neck dislocation and decapitation. Coronal 350 µm slices were cut using a vibrating microtome (Microm HM650V) in a pre-chilled cutting solution containing (in mM): 65 Sucrose, 85 NaCl, 2.5 KCl, 1.25 NaH_2_PO_4_, 7 MgCl_2_, 0.5 CaCl_2_, 25 NaHCO_3_ and 10 glucose, pH 7.2–7.4 and bubbled with carbogen (95% O_2_/5% CO_2_). Slices were then incubated for at least 1 h in a storage chamber containing artificial cerebrospinal fluid (aCSF; in mM): 130 NaCl, 3.5 KCl, 1.2 NaH_2_PO_4_, 1 MgCl_2_, 1.5 CaCl_2_, 24 NaHCO_3_ and 10 glucose, pH 7.2–7.4, at RT and bubbled with carbogen. When required, slices were transferred to a recording chamber superfused with aCSF, bubbled with carbogen (30 ^о^C and perfusion speed of 2 ml/min). Pharmacological manipulations were delivered by bath application of drugs through the perfusion system for at least 15 minutes. Stock solutions were generated, aliquoted, and stored at -20^о^C. On an experiment day, stock solution was added to the aCSF to achieve the desired final concentration (in µM): 10 bumetanide (NKCCI inhibitor), 10 VU0463271 (KCC2 inhibitor). All drugs were purchased from Tocris Bioscience.

### Patch clamp recordings

Patch pipettes were pulled from standard wall borosilicate glass capillaries (2-5 MΩ). For gramicidin perforated patch recording pipettes were filled with a high KCl based internal solution to be able to monitor the integrity of the perforated patch, containing (in mM): 135 KCl, 4 Na_2_ATP, 0.3 Na_3_GTP, 2 MgCl_2_, and 10 HEPES. Osmolarity was adjusted to 290 mOsM and the pH was adjusted to 7.35 with KOH. Gramicidin (Calbiochem) was dissolved in dimethylsulfoxide (DMSO) to achieve a stock concentration of 4 mg/ml. This was then diluted into the internal solution on the day of the experiment to achieve a final concentration of 80 μg/ml. The resulting solution was vortexed for 40 s, sonicated for 10 s, then filtered through a 0.45 μm pore cellulose acetate membrane filter (Nalgene) and used immediately.

Neurons were visualized under a 60x water-immersion objective (Olympus BX51WI). Recordings were performed with an Axopatch 1D amplifier (Molecular Devices), acquired using WinWCP Strathclyde software (V.3.9.7; University of Strathclyde) and stored for off-line analysis. Recordings were made when the series resistance had stabilized to approximately 100 MΩ (approximately 30 min after gigaseal formation). [Cl^-^]_i_ measurements were performed by activating either GABA_A_ or glycine receptors by delivering short ‘puffs’ of GABA (Tocris Bioscience, 100 μM) via a patch pipette placed in the vicinity of the cell soma and connected to a picospritzer (5–10 psi for 20– 40 ms; General Valve). Puffs were delivered at a low frequency (15 s intervals) to ensure recovery of chloride homeostasis. For whole-cell patch clamp recordings, pipettes were filled with a cesium-gluconate based internal solution containing (in mM): 115 CsMeSO_3_, 4 Na_2_ATP, 0.4 Na_3_GTP, 10 Na2-phosphocreatine, 5 TEACl, 2 MgCl_2_, and 10 HEPES, 10 EGTA. Osmolarity was adjusted to 290 mOsM and the pH was adjusted to 7.35 with KOH and the series resistance was below 40 MΩ.

### E_GABAA_ measurement

EGABA_A_ measurements were performed in voltage clamp mode from a holding potential of -70 mV. Test voltage ramps (a saw-tooth, down-up function of 500 ms duration, with a minimum of -90 mV and maximum of -50 mV) were delivered at baseline (control ramp) and near the peak of the GABA-evoked current (GABA ramp). For each neuron, I-V curves of the control and GABA ramps were generated using linear fits after 70% series resistance correction. EGABA_A_ was defined as the membrane potential at which the control and GABA ramp currents intersected, which was equivalent to the membrane potential at which the difference between the GABA and control ramp currents was equal to zero. EGABA_A_ for each data point was a mean of 5-10 measurements.

### Synaptic stimulation and LTP induction

Synaptic inputs were activated via a sharpened epoxy-insulated tungsten stimulating electrode (A-M Systems; tip <10 μm) placed in lower L2/3. Pulses were delivered by an isolated voltage stimulator (Digitimer, DS2), controlled via TTL pulses from the electrophysiology software. In LTP experiments, the peak amplitude of the subthreshold monosynaptic EPSP (less than 10 mV) was monitored for a 5-10 minute baseline period at a frequency of 0.1 Hz. LTP induction consisted of 100 stimulus trains (each comprising four stimuli at 100 Hz, every 10 s), using the lowest stimulus intensity required to elicit one spike per stimulus. Following LTP induction, responses were monitored using the same subthreshold stimulus intensity as before LTP induction, for a period of at least 20 minutes at a frequency of 0.1 Hz.

### Data analysis and statistics

All data analysis was performed in Matlab, while all statistical tests were performed using InStat (Graphad). No statistical methods were used to pre-determine sample sizes, but our sample sizes are similar to those reported in previous publications^36,50–53^. Data was assessed for normality using a Kolmogorov-Smirnov test, and then the appropriate parametric or non-parametric test was performed. Appropriate correction such as the Welch’s correction were applied when comparing data with unequal standard deviations. All tests were two-tailed, with a confidence level of at least a 95%. Effect size was estimated using Cohen’s D test, computed using the ‘computeCohen_d’ function in Matlab and was denoted as ‘d’. Unless stated otherwise, values in the text represent mean ± stdev. No animals or data points were excluded from the analyses.

## References

1. Tononi, G. & Cirelli, C. Sleep and the Price of Plasticity: From Synaptic and Cellular Homeostasis to Memory Consolidation and Integration. Neuron vol. 81 12–34 (2014).

2. Klinzing, J. G., Niethard, N. & Born, J. Mechanisms of systems memory consolidation during sleep. Nature Neuroscience vol. 22 1598–1610 (2019).

3. De Vivo, L. et al. Ultrastructural evidence for synaptic scaling across the wake/sleep cycle. Science 355, 507–510 (2017).

4. Vyazovskiy, V. V., Cirelli, C., Pfister-Genskow, M., Faraguna, U. & Tononi, G. Molecular and electrophysiological evidence for net synaptic potentiation in wake and depression in sleep. Nat. Neurosci. 11, 200–208 (2008).

5. Zhou, Y. et al. REM sleep promotes experience-dependent dendritic spine elimination in the mouse cortex. Nat. Commun. 11, 4819 (2020).

6. Aton, S. J. et al. Mechanisms of Sleep-Dependent Consolidation of Cortical Plasticity. Neuron 61, 454–466 (2009).

7. Brüning, F. et al. Sleep-wake cycles drive daily dynamics of synaptic phosphorylation. Science 366, (2019).

8. Chauvette, S., Seigneur, J. & Timofeev, I. Sleep Oscillations in the Thalamocortical System Induce Long-Term Neuronal Plasticity. Neuron 75, 1105–1113 (2012).

9. Aton, S. J., Suresh, A., Broussard, C. & Frank, M. G. Sleep Promotes Cortical Response Potentiation Following Visual Experience. Sleep 37, 1163–1170 (2014).

10. González-Rueda, A., Pedrosa, V., Feord, R. C., Clopath, C. & Paulsen, O. Activity-Dependent Downscaling of Subthreshold Synaptic Inputs during Slow-Wave-Sleep-like Activity In Vivo. Neuron 97, 1244–1252.e5 (2018).

11. Goode, L. K. et al. Examination of Diurnal Variation and Sex Differences in Hippocampal Neurophysiology and Spatial Memory. eneuro 9, ENEURO.0124-22.2022 (2022).

12. Chaudhury, D., Wang, L. M. & Colwell, C. S. Circadian Regulation of Hippocampal Long-Term Potentiation. J. Biol. Rhythms 20, 225–236 (2005).

13. Liu, Z. W., Faraguna, U., Cirelli, C., Tononi, G. & Gao, X. B. Direct evidence for wake-related increases and sleep-related decreases in synaptic strength in rodent cortex. J. Neurosci. 30, 8671–8675 (2010).

14. Hengen, K. B., Torrado Pacheco, A., McGregor, J. N., Van Hooser, S. D. & Turrigiano, G. G. Neuronal Firing Rate Homeostasis Is Inhibited by Sleep and Promoted by Wake. Cell 165, 180–191 (2016).

15. Torrado Pacheco, A., Bottorff, J., Gao, Y. & Turrigiano, G. G. Sleep Promotes Downward Firing Rate Homeostasis. Neuron 109, 530–544.e6 (2021).

16. Tononi, G. & Cirelli, C. Sleep and synaptic homeostasis: A hypothesis. Brain Res. Bull. 62, 143–150 (2003).

17. Seibt, J. & Frank, M. G. Primed to Sleep: The Dynamics of Synaptic Plasticity Across Brain States. Frontiers in Systems Neuroscience vol. 13 (2019).

18. Alfonsa, H. et al. Intracellular chloride regulation mediates local sleep pressure in the cortex. Nat. Neurosci. (2022) doi:10.1038/s41593-022-01214-2.

19. Doyon, N., Vinay, L., Prescott, S. A. & De Koninck, Y. Chloride Regulation: A Dynamic Equilibrium Crucial for Synaptic Inhibition. Neuron 89, 1157–1172 (2016).

20. Blaesse, P., Airaksinen, M. S., Rivera, C. & Kaila, K. Cation-Chloride Cotransporters and Neuronal Function. Neuron vol. 61 820–838 (2009).

21. Düsterwald, K. M. et al. Biophysical models reveal the relative importance of transporter proteins and impermeant anions in chloride homeostasis. Elife 7, 39575 (2018).

22. Isaacson, J. S. & Scanziani, M. How Inhibition Shapes Cortical Activity. Neuron 72, 231–243 (2011).

23. Okun, M. & Lampl, I. Instantaneous correlation of excitation and inhibition during ongoing and sensory-evoked activities. Nat. Neurosci. 11, 535–537 (2008).

24. Pouille, F. & Scanziani, M. Enforcement of temporal fidelity in pyramidal cells by somatic feed-forward inhibition. Science 293, 1159–1163 (2001).

25. Akerman, C. J. & Cline, H. T. Depolarizing GABAergic Conductances Regulate the Balance of Excitation to Inhibition in the Developing Retinotectal Circuit In Vivo. J. Neurosci. 26, 5117 LP – 5130 (2006).

26. Jedlicka, P., Deller, T., Gutkin, B. S. & Backus, K. H. Activity-dependent intracellular chloride accumulation and diffusion controls GABAA receptor-mediated synaptic transmission. Hippocampus 21, 885–898 (2011).

27. Mapelli, J., Gandolfi, D., Vilella, A., Zoli, M. & Bigiani, A. Heterosynaptic GABAergic plasticity bidirectionally driven by the activity of pre- and postsynaptic NMDA receptors. Proc. Natl. Acad. Sci. 113, 9898–9903 (2016).

28. Sjöström, P. J., Turrigiano, G. G. & Nelson, S. B. Rate, Timing, and Cooperativity Jointly Determine Cortical Synaptic Plasticity. Neuron 32, 1149– 1164 (2001).

29. Lisman, J. & Spruston, N. Postsynaptic depolarization requirements for LTP and LTD: a critique of spike timing-dependent plasticity. Nat. Neurosci. 8, 839– 841 (2005).

30. Meissner-Bernard, C., Tsai, M. C., Logiaco, L. & Gerstner, W. Dendritic Voltage Recordings Explain Paradoxical Synaptic Plasticity: A Modeling Study. Frontiers in Synaptic Neuroscience vol. 12 (2020).

31. Stuart, G. J. Determinants of Spike Timing-Dependent Synaptic Plasticity. Neuron 32, 966–968 (2001).

32. Clopath, C., Büsing, L., Vasilaki, E. & Gerstner, W. Connectivity reflects coding: a model of voltage-based STDP with homeostasis. Nat. Neurosci. 13, 344–352 (2010).

33. Richardson, G. S., Moore-Ede, M. C., Czeisler, C. A. & Dement, W. C. Circadian rhythms of sleep and wakefulness in mice: analysis using long-term automated recording of sleep. Am. J. Physiol. Integr. Comp. Physiol. 248, R320–R330 (1985).

34. Kyrozis, A. & Reichling, D. B. Perforated-patch recording with gramicidin avoids artifactual changes in intracellular chloride concentration. J. Neurosci. Methods 57, 27–35 (1995).

35. Huberfeld, G. et al. Perturbed Chloride Homeostasis and GABAergic Signaling in Human Temporal Lobe Epilepsy. J. Neurosci. 27, 9866 LP – 9873 (2007).

36. Choi, H. J. et al. Excitatory actions of GABA in the suprachiasmatic nucleus. J. Neurosci. 28, 5450–5459 (2008).

37. Sivakumaran, S. et al. Selective Inhibition of KCC2 Leads to Hyperexcitability and Epileptiform Discharges in Hippocampal Slices and In Vivo. J. Neurosci. 35, 8291 LP – 8296 (2015).

38. Pracucci, E., et al. Circadian rhythm in cortical chloride homeostasis underpins variation in network excitability. bioRxiv 2021.05.12.443725 (2021) doi:10.1101/2021.05.12.443725.

39. Noya, S. B. et al. The forebrain synaptic transcriptome is organized by clocks but its proteome is driven by sleep. Science 366, (2019).

40. Bridi, M. C. D. et al. Daily Oscillation of the Excitation-Inhibition Balance in Visual Cortical Circuits. Neuron 105, 621–629.e4 (2020).

41. Zong, F.-J. et al. Circadian time- and sleep-dependent modulation of cortical parvalbumin-positive inhibitory neurons. EMBO J. 42, e111304 (2023).

42. Bliss, T. V. P. & Collingridge, G. L. A synaptic model of memory: long-term potentiation in the hippocampus. Nature 361, 31–39 (1993).

43. Ascher, P. & Nowak, L. The role of divalent cations in the N-methyl-D-aspartate responses of mouse central neurones in culture. J. Physiol. 399, 247–266 (1988).

44. Collingridge, G. L., Herron, C. E. & Lester, R. A. Synaptic activation of N-methyl-D-aspartate receptors in the Schaffer collateral-commissural pathway of rat hippocampus. J. Physiol. 399, 283–300 (1988).

45. Bi, G. & Poo, M. Synaptic Modification by Correlated Activity: Hebb’s Postulate Revisited. Annu. Rev. Neurosci. 24, 139–166 (2001).

46. Ben-Ari, Y. Excitatory actions of gaba during development: the nature of the nurture. Nat. Rev. Neurosci. 3, 728–739 (2002).

47. Sjöström, P. J. & Häusser, M. A Cooperative Switch Determines the Sign of Synaptic Plasticity in Distal Dendrites of Neocortical Pyramidal Neurons. Neuron 51, 227–238 (2006).

48. Aton, S. J., Suresh, A., Broussard, C. & Frank, M. G. Sleep Promotes Cortical Response Potentiation Following Visual Experience. Sleep 37, 1163–1170 (2014).

49. Diering, G. H. et al. Homer1a drives homeostatic scaling-down of excitatory synapses during sleep. Science 355, 511–515 (2017).

50. Alfonsa, H. et al. The Contribution of Raised Intraneuronal Chloride to Epileptic Network Activity. J. Neurosci. 35, 7715–7726 (2015).

51. Huber, R., Deboer, T. & Tobler, I. Topography of EEG Dynamics After Sleep Deprivation in Mice. J. Neurophysiol. 84, 1888–1893 (2000).

52. Palchykova, S., Winsky-Sommerer, R., Meerlo, P., Dürr, R. & Tobler, I. Sleep deprivation impairs object recognition in mice. Neurobiol. Learn. Mem. 85, 263–271 (2006).

53. Krone, L. B. et al. A role for the cortex in sleep–wake regulation. Nat. Neurosci. 2021 249 24, 1210–1215 (2021).

